# A hermetically closed sample chamber enables time-lapse nano-characterization of pathogenic microorganisms *in-vitro*

**DOI:** 10.1101/2025.01.08.631953

**Authors:** Esther Braun, Santiago H. Andany, Mustafa Kangül, Navid S. Asmari, John D. McKinney, Georg E. Fantner

## Abstract

Pathogenic microorganisms, such as pathogenic mycobacteria, pose a global health burden. Studying these organisms is crucial for gaining detailed knowledge about the pathogens and the diseases they cause. To handle pathogenic organisms, specific biosafety measures appropriate to the virulence of the organism must be fulfilled, most importantly ensuring that all manipulations of pathogenic material are performed within a confined environment. Atomic force microscopy (AFM) is a powerful technique to study biological samples at nanometer-scale resolution, yielding also mechanical properties, all while maintaining physiological conditions. However, standard AFM sample holders do not meet stringent biosafety requirements since they do not constitute a confined system. AFM imaging relies on direct contact between the cantilever and the sample and is sensitive to mechanical interference, rendering conventional containment systems for handling infectious substances inapplicable. Here, we introduce a hermetically sealed AFM sample chamber that meets biosafety demands while satisfying the mechanical and optical constraints of correlated optical microscopy and AFM. We imaged various pathogenic mycobacteria to demonstrate the chamber’s versatility and effectiveness in containing biohazardous materials. This sample chamber enables high-resolution, time-lapse correlated imaging and biomechanical characterization of pathogenic microorganisms *in-vitro*. It broadens the scope of research with pathogenic microorganisms under safe and controlled conditions.

## Introduction

Atomic force microscopy (AFM) is a powerful tool to study fine surface structures and subtle details of biological samples in the nanometer range^1,2^, that are invisible with conventional diffraction-limited optical microscopy. AFM is particularly well-suited for visualizing biological specimens with high-resolution^3^, while maintaining physiological conditions within liquid environments^4^. Using live-cell^5–9^, time-lapse AFM imaging^10–12^ of bacteria, allows to investigate dynamic biological processes unfolding over time *in-vitro*. Beyond the sample’s topography, AFM imaging also yields biomechanical properties^13–17^ and the cantilever can be used as a nanomanipulation tool^13,18^. Correlated imaging with fluorescence microscopy additionally enables the exploration of molecular phenomena^1,19,20^. These unique imaging capabilities of AFM have contributed to our current understanding of non-pathogenic mycobacteria^5,6^ and offer potential for advancing research on pathogenic microorganisms, by providing high-resolution information *in-vitro*. However, using pathogenic material with AFM poses additional challenges.

Pathogenic microorganisms are categorized into biosafety levels (BSLs) 1 to 4, based on their virulence and the potential biosafety risks. Depending on their biosafety level, pathogenic organisms must be handled with specific measures in dedicated BSL laboratory facilities. This includes additional requirements applied to technical instruments^21^. The principal biosafety regulations require performing all manipulations of pathogenic biological material within a controlled and enclosed environment. This implies the use of primary containments (i.e. first line of physical barriers preventing the release of hazardous biological agents) such as a biosafety cabinet (BSC)^21,22^. Standard AFM setups do not meet these biosafety requirements, because they are not a confined system. AFMs are scanning probe microscopes and rely on direct physical interactions between the probe and the sample. This renders conventional containment systems inapplicable. Those involve rigid barriers that prevent direct contact with the sample and obstruct the movement of the AFM scanner. Strong airflows and significant mechanical vibrations, which can induce mechanical disturbances leading to imaging artifacts, make using an AFM within a standard BSC unsuitable. Beniac et al.^23^ and Barlerin et al.^24^ introduced custom-built microscope biosafety cabinets for a scanning electron microscope (SEM) and a multiphoton microscope, respectively. These enclosures provide a sealed enclosure for the entire microscope under negative pressure, utilizing high-efficiency particulate air (HEPA) filters for air exchange. A glove box system facilitates access for microscope operation. Commercial glove box systems for AFM imaging are available and are typically used for the measurement of non-biological samples that require environmental control. This approach prevents the need for modifying the actual microscope but presents several challenges. Enclosing the complete microscope can lead to microscope components coming into contact with pathogenic material, resulting in a contamination of that part. Consequently, a decontamination procedure, such as using vaporized hydrogen peroxide, is necessary before repair or maintenance. This could damage sensitive electronic or optical parts. Using a glove box system also complicates sample handling and microscope operation.

Even though AFM studies involving BSL-2 classified bacteria have been described^25–29^, they do not indicate the use of AFM setups that fulfill the BSL requirements. A customized AFM system (Nanowizard 4, JPK-Bruker) with additional sealing of the AFM scanner^30^ has been utilized within a BSL-3 laboratory for research on both infectious and inactivated BSL-3 microorganisms^30–32^. However, when the microscope is used to study airborne pathogens, full hermetic sealing is required to prevent any release of pathogenic material, as these organisms are primarily transmitted through the inhalation of droplets. This is particularly crucial for pathogens where even a low pathogenic load can potentially cause infection^33^. Imaging such pathogenic micro-organisms is thus particularly challenging for AFM and calls for the development of a containment method that addresses both biosafety and AFM requirements. In this work, we present a hermetically sealed AFM sample chamber (Figure 1A) designed to fulfill the biosafety demands required for working with airborne BSL-2 and BSL-3 pathogens while accommodating the mechanical constraints of correlated optical microscope and AFM (Figure 1B), thereby facilitating time-lapse nano-characterization of pathogenic microorganisms *in-vitro*.

**Figure 1:**
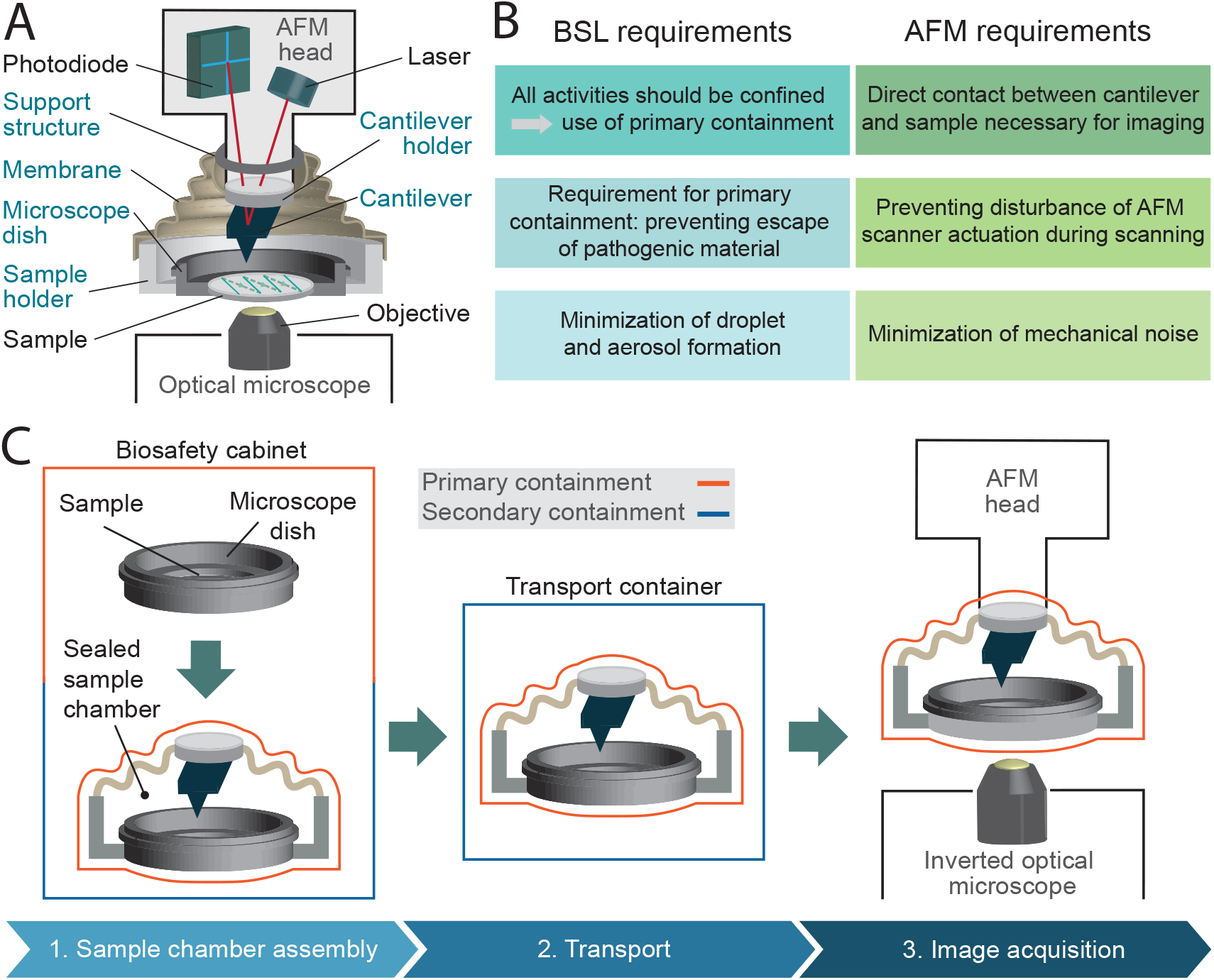
Concept of the hermetically sealed sample chamber. **A** Schematic of a combined optical microscope and AFM setup with the hermetically sealed sample chamber, which fully confines the pathogenic material. **B** List of primary biosafety and AFM requirements that form the basis for the design considerations of the hermetically sealed sample chamber. **C** Overview of the sample chamber handling approach, illustrating a series of controlled steps to ensure user safety. (1) All procedures involving direct manipulation of pathogenic material, including sample preparation, are initially performed within a biosafety cabinet. The entire sample preparation process, which includes immobilizing the sample and performing washing steps, is carried out within the microscope dish inside a BSC. Once the sample chamber is securely sealed and the pathogenic material completely confined, it is removed from the biosafety cabinet. (2) For transport to the AFM setup, the sample chamber is placed inside a secondary containment. (3) During AFM imaging, the sample chamber serves as the primary containment, maintaining the confinement of the pathogenic material. Throughout this process, multiple containment strategies are employed to ensure the secure handling of the pathogenic material.

## Main

### Design concept of the hermetically sealed sample chamber

To allow AFM imaging within a biosafety environment, we designed a hermetically closed sample chamber concept to completely seal fluids and vapors while preserving the AFM functionality. The concept includes three design approaches: We conceptualized the sample chamber as an independent sealed unit that includes the cantilever and cantilever holder, allowing it to be assembled separately from the AFM head with the temporal use of specific support structures. A membrane ensures airtight sealing and provides sufficient flexibility for the AFM scanner movement on two levels. We implemented these concepts by designing and manufacturing a sample chamber for a specific AFM head (Dimension 3100, Digital Instruments) and compatible with a standard microscope dish (PELCO® Clear Wall Glass Bottom Dish). We developed a specific handling procedure to guarantee a consistent confinement of biohazardous material in a closed environment (Figure 1C, Supplementary Figure 1). All open manipulations of pathogenic material, including the sample preparation process, are carried out inside a BSC. Once the sample preparation is completed, the microscope dish is inserted into the partly assembled sample chamber and sealed (Figure 1C, step 1). The sample chamber is then transferred out of the BSC and to the AFM setup within a second containment, capable of withstanding an unexpected fall without damage (Figure 1C, step 2). During the AFM imaging, the hermetically sealed sample chamber itself serves the central role of a primary containment (Figure 1C, step 3).

### Design choices to comply with biosafety requirements

We designed a completely hermetically sealed system to fulfill the main requirement of primary containments: the minimization (BSL-2) or the prevention (BSL-3) of escape of pathogenic material^22^. We achieve this by using multiple containment methods, including specifically designed silicone membranes and rubber O-rings, all integrated with different clamping mechanisms (Figure 2A-B, Supplementary Figure 2, Supplementary Figure 3). Conventionally, the cantilever holder is first attached to the AFM scanner and subsequently brought into contact with the sample. Our sample chamber design, as an independent sealed unit encompassing the cantilever holder, eliminates the necessity to introduce the AFM head into the BSC during the assembly process for imaging preparation. This mitigates the risk of contaminating the AFM head, whose intricate electronic and optical components are difficult to decontaminate. To facilitate the attachment of the cantilever holder, as an integrated part of the sample chamber, to the AFM scanner, we designed a custom cantilever holder featuring a magnetic mounting system. The magnets provide magnetically assisted semi-automatic mounting and secure fixation, while a dowel pin and sleeve mechanism ensures alignment (Figure 2A, Supplementary Figure 4). Further primary safety requirements include minimizing aerosol and droplet formation, eliminating sharp edges and unguarded moving parts, and ensuring the ability to decontaminate all materials^21,22^. To meet these criteria, we designed the sample chamber with an airtight membrane to contain aerosols and droplets during AFM imaging in liquid. The chamber is free of sharp edges, and all moving parts are securely enclosed by 3D-printed housings to prevent damaging personal safety equipment. Additionally, all components are constructed from materials that can resist double decontamination treatment by both chemical decontamination and steam autoclaving (Supplementary Table 1). We chose a sample chamber design integrating a disposable microscope dish to further minimize the risk of contamination by reducing the number of sample chamber components in contact with biohazardous material. This allows the entire sample preparation process to be carried out solely using the microscope dish, without the need to mount it into the sample chamber beforehand. Especially during sample preparation, tasks such as sample immobilization and multiple washing steps carry an elevated risk of potential spillage and contamination of sample chamber components. Enabling a sample preparation process independent of the sample chamber reduces the risk of contaminations, while offering greater flexibility and easier handling of the microscope dish compared to managing the entire sample chamber assembly (Supplementary Figure 1).

**Figure 2:**
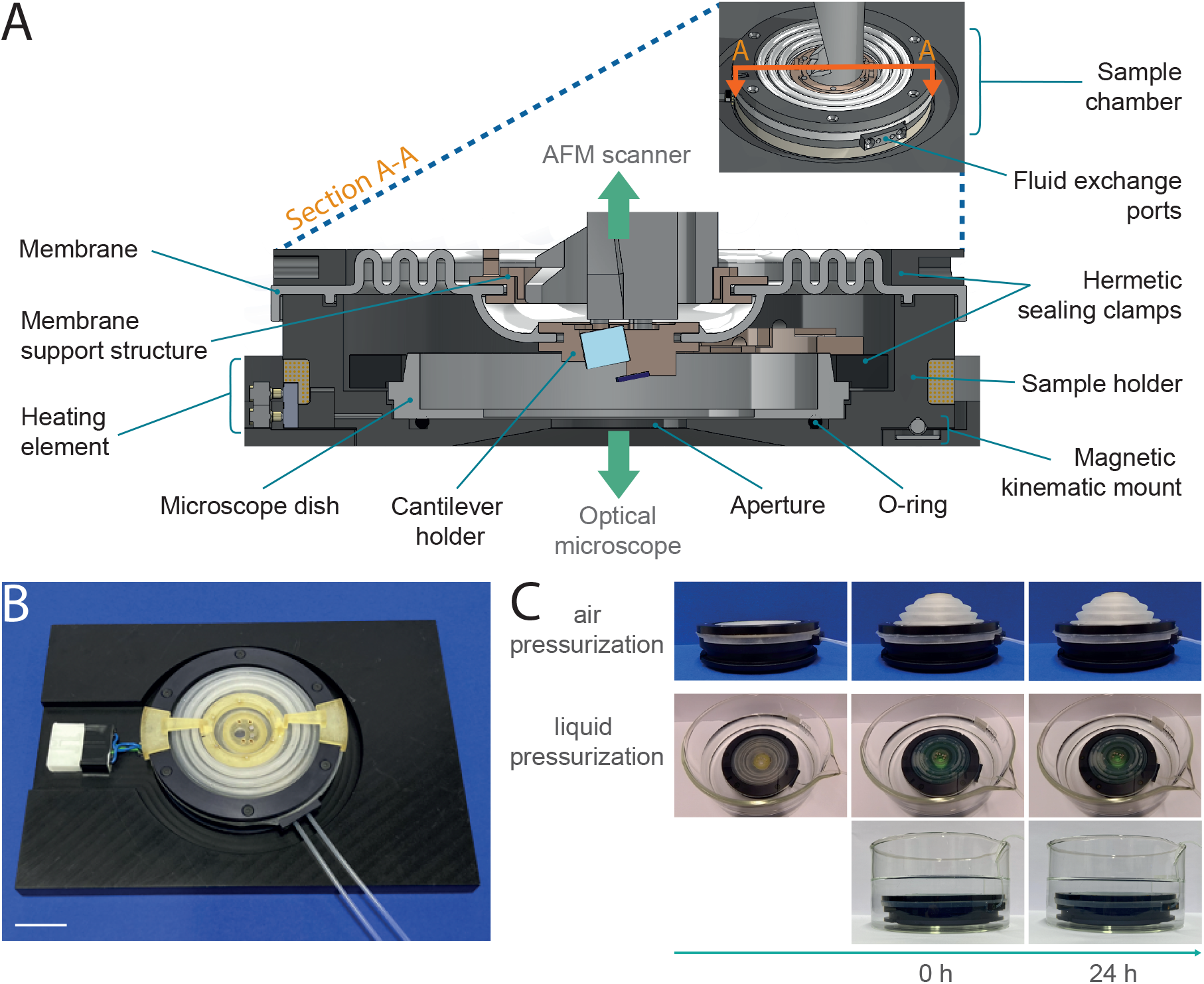
Design of sample chamber and evaluation of biosafety compliance. **A** Cross-section (A-A) of the sample chamber showing its main components. The sample chamber is positioned above an inverted microscope, while the cantilever holder is connected to the AFM scanner placed on top of the sample chamber. **B** Image of the fully sealed sample chamber assembly, featuring tubes for fluid exchange and a connector for temperature control. The chamber is mounted in a metal frame to enhance mechanical stability (scale bar: 20 mm). **C** Evaluation of the sample chamber’s airtightness and leakproofness. For airtightness testing, the chamber was pressurized with air and monitored for any deflation over a 24-hour period (top). To assess leakproofness, the chamber was pressurized with a dyed liquid and submerged in a transparent liquid (bottom).

### Design choices to maintain AFM performance

We ensure undeteriorated AFM performance by preventing scanner movement disturbances and minimizing additional mechanical noise. The membrane provides sufficient flexibility on two levels. The outer part of the membrane allows coarse x-y-z movement of the sample chamber with respect to the AFM scanner housing. The inner part of the membrane provides flexibility in x-y-z direction for fine AFM scanner movement (Figure 2A, Supplementary Figure 5A). To ensure adequate sealing, the cantilever holder must be in direct contact with the membrane which can disturb the scanner actuation. To minimize interference with the scanner motion, we integrated a membrane support structure system with dual use. It includes a support structure fixed to the membrane and anchored to the AFM scanner enclosure. The support structure reduces the membrane’s weight on the cantilever holder, minimizing distortions in piezo-actuator movement and decreasing coupling effects. Additionally, temporary support arms can be attached via a slot-and-tab mount to prevent the cantilever holder from descending during assembly and transport, protecting the cantilever from damage (Figure 2A, Supplementary Figure 4B-C, Supplementary Figure 5B).

### Enhanced imaging capabilities with additional sample chamber features

An aperture at the bottom of the sample holder allows correlated AFM and optical imaging with an inverted light microscope (Supplementary Figure 6A, Supplementary Figure 7). We use a glass-bottom microscope dish (PELCO^®^ Clear Wall Glass Bottom Dish) mounted in the sample holder for use with high-NA oil-immersion objectives. Temperature and fluid control systems allow *in-vitro* time-lapse imaging under physiologically relevant conditions (Figure 2A-B, Supplementary Figure 6). An integrated heating element and a temperature sensor enable stable regulation of the temperature setpoint. The sample holder has two sealed ports, each with an inlet and outlet for medium exchange. To handle the potentially contaminated medium from the fluid outlet, we incorporated a waste deactivation system that directs the waste media into a vessel containing disinfectant.

### Sample chamber complies with BSL 2 and BSL 3 requirements

We then tested the sealing of the closed sample chamber to ensure biosafety compliance by performing airtightness and liquid leakproof tests (Figure 2C, Supplementary Figure 8, Supplementary Videos 1-5). To verify airtightness, we pressurized the sample chamber with air and monitored for deflation over a 24-hour period. The absence of deflation indicated effective airtight sealing. We repeated the airtightness test over 100 cycles with shorter test periods, to simulate a multitude of imaging sessions. To assess the endurance of the silicone membrane, we simulated an aging process by repeatedly autoclaving the membrane and subsequently testing the airtightness (Supplementary

Table 2). For leakproof assessment, we pressurized the sample chamber with a dyed liquid and then submerged it in a transparent liquid. Transparency remaining after 24 hours indicated no leakage. These assessments confirmed airtightness and liquid leakproofness, even after multiple imaging sessions and aging of the membrane. Additionally, we demonstrated that the sample chamber components are resistant to steam autoclaving, the preferred method of decontamination^21^, and chemical disinfectants with microbiological efficacy against commonly studied pathogenic organisms (Supplementary Table 1). Based on these tests, the sample chamber received official BSL-2 and BSL-3 authorization from the *Swiss Federal Office for Public Health*, verifying the compliance with the biosafety requirements.

### Applying the sample holder to image pathogenic mycobacteria

To showcase the versatility and imaging capabilities of the sample chamber, we performed multiple proof-of-concept experiments. We used it to image different types of pathogenic mycobacteria across various imaging modalities. We imaged both the smooth (Figure 3A) and rough (Figure 3B) variants of *M. abscessus* and *Mycobacterium marinum* (Figure 3C), which are BSL-2 classified pathogens. *M. abscessus* is among the most virulent and drug-resistant rapidly growing nontuberculous mycobacteria and still lacks effective antibiotic treatments^34,35^. *M. marinum* is a slow-growing mycobacterium typically contracted through a soft tissue injury in either freshwater or saltwater environments^34^. The sample chamber allows for the acquisition of biomechanical cell properties through multiple AFM image channels, such as stiffness, adhesion, and dissipation (Figure 3C). It also supports combined optical and AFM imaging. We obtained correlated fluorescence and AFM data of *M. abscessus* (smooth variant) stained with BODIPY™ FL Vancomycin (Figure 3D, Supplementary Figure 10) and a green fluorescent protein (GFP)-expressing *M. marinum* strain (Figure 3E). Moreover, the sample chamber enables live-cell time-lapse imaging. We performed time-lapse AFM imaging of *M. abscessus* (smooth variant) over more than 9 hours (Figure 3F).

**Figure 3:**
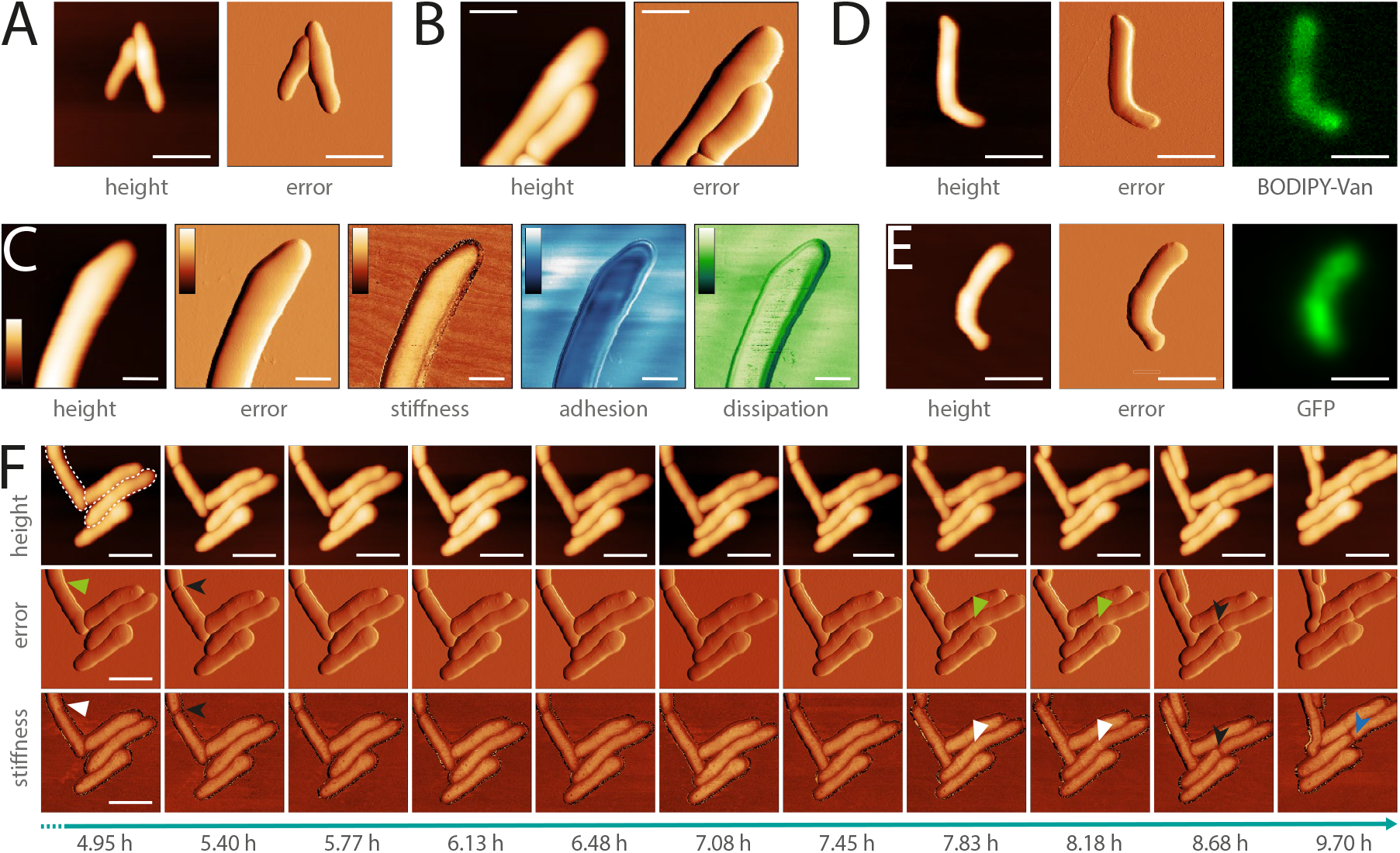
The sample chamber can be used for a range of pathogenic mycobacteria (BSL-2) and for different imaging modalities. **A** AFM image of *M. abscessus* smooth variant (scale bar 2µm). **B** AFM image of *M. abscessus* rough variant (scale bar 1 µm). **C** Different channels (height, error, stiffness, adhesion, dissipation) of an AFM image of *M. marinum* (scale bar 1µm, color scale: height 0-750 nm, error 1.05-3.30 nN, stiffness (estimated by approach slope) 0.17-0.26, adhesion 0-0.25 nN, dissipation 0-0.27 meV). **D** Correlated optical and AFM image of *M. abscessus* smooth variant stained with BODIPY™ FL Vancomycin (scale bar 2µm). **E** Correlated optical and AFM image of *M. marinum* GFP (scale bar 2µm). The bottom cell tip was not completely immobilized on the surface and was slightly moving during the AFM imaging. **F** Timelapse AFM images over a period >9 h of *M. abscessus* smooth variant cells. Two cells (dotted outline) show appearance of the pre-cleavage furrow (green arrowhead) with associated increased stiffness (white arrowhead). After cell cleavage (black arrowhead) a division scar can be observed (blue arrowhead) (scale bar 2µm).

Next, we studied morphological events during cell division in *M. abscessus*, by performing a high-resolution topographic and nanomechanical characterization *in-vitro*. Previous AFM studies of the related non-pathogenic bacterium *Mycobacterium smegmatis* showed that a pre-cleavage furrow (PCF), observable as an indentation in the cell wall, forms at the future division site, presenting the initial morphological characteristic of a developing division event^10^. The PCF is associated with a stress concentration, observable as a higher level of stiffness, which increases over time cumulating in cell cleavage^13^. These events are involved in the mechanical mechanism underlying mycobacterial cell division^13^. In order to better understand cell division in *M. abscessus*, we investigated whether these events also occur in this species. We did this by performing time-lapse AFM imaging while simultaneously acquiring biomechanical properties of the M. abscessus smooth variant (Figure 4, Supplementary Figure 11). Similar to *M. smegmatis*, we observed the formation of a PCF and an increased stiffness at the surface area surrounding the PCF in *M. abscessus* (Figure 4A, Supplementary Figure 10B). When following single cells over time, we observed that the stiffness at the PCF increases with cell cycle progression and cell cleavage occurs at the PCF (Figure 4B, Figure 3F, Supplementary Figure 11A, C, D). We also observed ‘division scars’ after cell cleavage, which are a characteristic phenotype of the mycobacterial cell division process^36,37^ (Supplementary Figure 11D). This suggests that the cell division process in *M. abscessus* involves similar morphological events as those observed in non-pathogenic mycobacteria.

**Figure 4:**
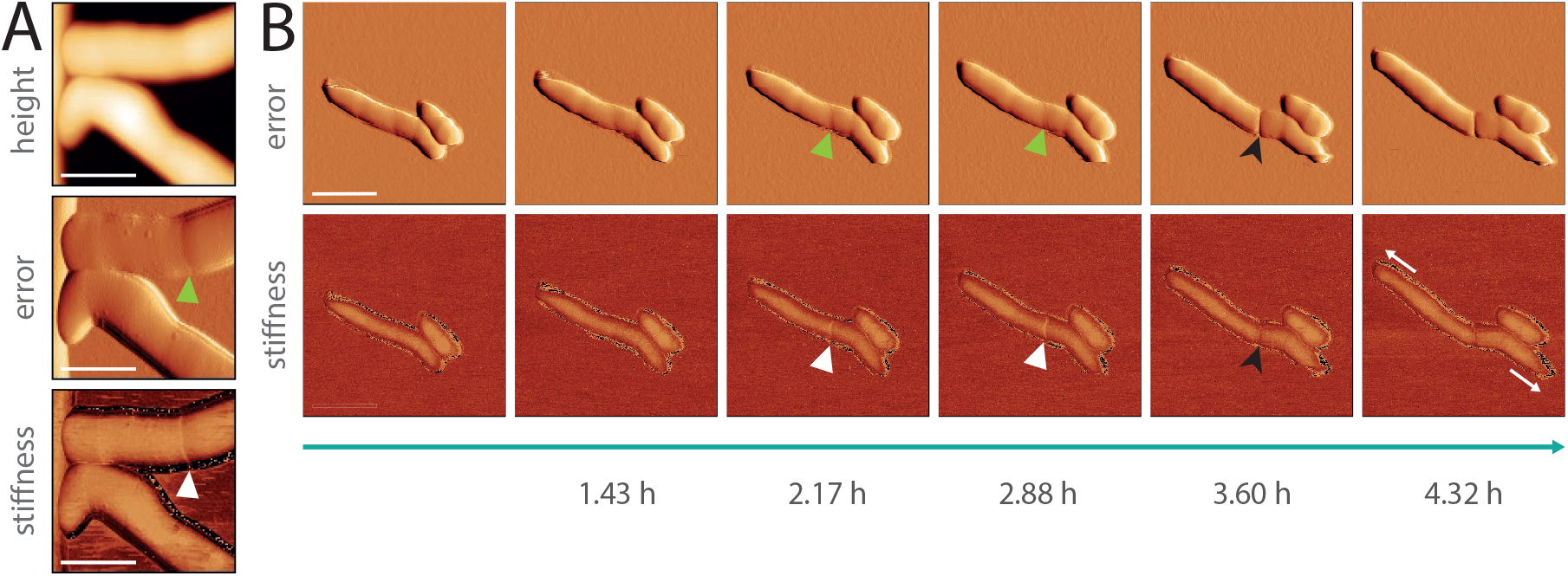
Characterization of cell division events of pathogenic mycobacteria *in-vivo*(*M. abscessus* smooth variant) **A** AFM images of cells before cell cleavage. The error image (middle) shows the formation of the pre-cleavage furrow (green arrowhead) and the stiffness image (bottom) an increased stiffness level at the area surrounding the pre-cleavage furrow (white arrowhead) (scale bar 1µm). **B** Timelapse AFM images showing the appearance of the pre-cleavage furrow (green arrowhead) with associated increased stiffness (white arrowhead). The level of stiffness increases over time cumulating in cell cleavage at the position of the pre-cleavage furrow (black arrowhead). The cells are showing polar cell growth (white arrows) (scale bar 2µm).

## Discussion

Here, we present the concept and implementation of a hermetically sealed sample chamber that allows correlated optical and AFM time-lapse imaging of pathogenic organisms *in-vitro*. It permits to study pathogens under safe and controlled conditions, thereby expanding the potential of AFM in microbiological research. We demonstrate that our hermetically sealed sample chamber meets the dual requirements of biosafety containment and AFM imaging (Figure 2C, Supplementary Figure 9). We used the sample chamber to image multiple pathogenic mycobacteria using different imaging modalities and studied the cell cycle of *M. abscessus* through time-lapse AFM imaging and biomechanical characterization, revealing similar morphological cell division events to those observed in non-pathogenic mycobacteria. This showcased the chamber’s versatility for imaging pathogenic mycobacteria and acquiring biomechanical cell properties.

The containment strategy does not compromise AFM functionality but we identified two factors affecting the AFM performance: The mechanically unsupported area (aperture size) of the sample holder and the liquid volume (growth media volume) used during imaging adversely affect the system’s bandwidth (Supplementary Information, Supplementary Figure 9). They limit the scan rate because they create dynamics that interfere with the feedback loop by reducing the phase and gain margins. Additionally, an aperture decreases the structural rigidity of the microscope dish and increases susceptibility to mechanical vibrations. To partly address these issues, we used a thin layer of vacuum grease between the sample holder and microscope dish to maximize the support of the microscope dish at the area around the aperture. A larger media volume inside the microscope dish increases the overall actuated mass during scanner actuation and increases the coupling between the piezo actuator and the microscope dish. Despite posing challenges, an aperture is essential for correlated optical imaging and adequate liquid volume is imperative to nourish cells during *in-vitro* imaging. We identified a trade-off for both aperture size and media volume that enables performant correlated imaging *in-vitro*.

Our design concept integrates three key enabling concepts: (1) a sample chamber including the cantilever holder to form a pre-assembled sealed unit, (2) a membrane offering two-level flexibility for AFM scanner movement, and (3) an assembly support system for assembly independent from the AFM head. We implemented our sample chamber concept for a Dimension 3100 AFM scan head but the concept can be used to design sample chambers suitable for other AFMs. The sample chamber can also be customized for specific experimental requirements, for instance by using the second port as CO2 supply for imaging of mammalian cells.

Here, we tested the sample chamber with BSL-2 classified mycobacteria. The official authorization from the Swiss Federal Office for Public Health validates that the sample chamber also meets the stringent requirements for usage with high-risk airborne BSL-3 pathogens, such as Mycobacterium tuberculosis. By adapting the microscope dish surface functionalization, the sample chamber can also be used for studying Gram-negative bacteria, such as uropathogenic *Escherichia coli* (UPEC), *Vibrio cholerae* or *Pseudomonas aeruginosa*.

In conclusion, the development and implementation of a hermetically sealed AFM sample chamber open up several promising avenues for future research. The design of our sample chamber allows for high-resolution time-lapse imaging and biomechanical studies of pathogenic microorganism *in-vitro* under controlled biosafety conditions. This advancement will empower comprehensive studies of pathogenic microorganisms, ultimately contributing to a better understanding of pathogens.

## Material and methods

### Sample chamber design, fabrication and handling

The sample holder (7075 aluminium alloy, anodized) is designed to hold a commercial disposable microscope dish (PELCO® Clear Wall Glass Bottom Dishes, Ted Pella Inc., 50 x 7mm, glass 30mm, 14024/5) (Supplementary Figure 1, Supplementary Figure 2).

The sealing of the sample chamber (Supplementary Figure 3) is ensured with a specifically designed silicone membrane (SF13 RTV-2 silicone), rubber O-rings (FPM-75) and metal clamps (7075 aluminium alloy, anodized). The bottom interface of the sample holder is sealed using a rubber O-ring (Dichtelemente arcus, 117958, FPM-75, 42 x 1.2 mm) combined with vacuum grease, both positioned beneath the microscope dish, which is secured by a clamp. The top of the sample holder is sealed with a membrane, clamped at the outer edge to the sample holder and at the inner edge to the cantilever holder using an interference fit (Supplementary Figure 3).

The customized cantilever holder (3D printed, PC-like translucent, Accura 5530) includes a glass window (custom cut, Edmund optics, 25 x 4 mm, VIS-EXT coated, λ/4 N-BK7 window, 13-298), two disc magnets (NdFeB, 2 x 1 mm) to facilitate mounting it to the AFM scanner, and four pin sleeves (7305-0-15-15-47-27-10-0, Mill-Max Manufacturing Corp.) for registration (Supplementary Figure 4A). The membrane is attached to the cantilever holder via an interference fit in a groove in the cantilever holder (Supplementary Figure 4B). Additionally, the cantilever holder is constructed without any holes, ensuring a completely sealed system once attached to the membrane. The glass window is glued into the cantilever holder with epoxy (9200 structural epoxy, MG Chemicals) while ensuring that all interfaces are closed. The cantilever is mounted to the cantilever holder with wax.

The membrane support structure system (3D printed, PC-like translucent, Accura 5530) consists of three sub-components with a twofold purpose (Supplementary Figure 4B). A circular support structure (1) attaches to the membrane by an interference fit and holds a substantial part of the membrane weight via an additional anchor point (2) permanently fixed to the AFM scanner enclosure. Both parts are fixed together by a magnetic mount (NdFeB, 2 x 1mm, 1 x 1mm). This support structure minimizes distortions in the piezo-actuator movement by reducing the weight of the membrane attached to the cantilever holder. It also decreases the coupling effects from the membrane onto the cantilever holder. Additionally, support arms (3) can be temporarily attached to the membrane support structure by a slot-and-tab mount to facilitate the assembly and transport of the sample chamber while preventing the cantilever holder from descending and thereby safeguarding the fragile cantilever from potential damage (Supplementary Figure 4C).

The principal membrane concept involves flexibility on two levels: The outer section of the membrane enables x-y-z movement of the sample chamber relative to the AFM scanner housing (i.e., coarse z-approach, adjustment region of interest). The inner section allows x-y-z flexibility for the AFM scanner’s motion within the housing (i.e., z-topography feedback, line scanning during image acquisition). The membrane is fabricated by silicone molding with a 3D-printed custom-designed mold (Supplementary Figure 5A). In short, the addition-curing silicone rubber (SF13 RTV-2, Silikonfabrik) was mixed 1:1, air-bubbles were removed by degassing under vacuum, the mixed silicone was injected into the assembled mold, baked at 50° C for 1h and then released from the mold. The mold consists of several air-release cutouts to ensure an air-bubble-free membrane. Avoiding air bubbles is crucial, as they can create small holes in the membrane, compromising the airtightness of the sealed sample holder. Protruding parts from the air-bubble release holes were removed with a sharp razor plate.

During the imaging, the sample chamber is placed in a metal frame (6082 aluminium alloy, anodized) (Figure 2B, Supplementary Figure 3A). To increase the mechanical stability and to ensure the positioning of the sample chamber assembly, a magnetic kinematic mount is implemented at the interface between the sample holder and the sample holder frame (Figure 2A, Supplementary Figure 6A).

The closed sample chamber comprises several features to expand the imaging capabilities. Correlated AFM and optical imaging (inverted light microscope) are enabled by an aperture (10 mm) at the bottom of the sample holder (Supplementary Figure 6A). The chosen type of microscope dish that has a glass coverslip bottom (#1.5) and the sample holder design (i.e. aperture diameter and angle) permit the use of high-magnification oil-immersed objectives. The sample chamber heating (Supplementary Figure 6) is implemented by a heating element (isolated resistance wires, RD100/0.2, 15.6 Ω/m, Block) glued (heat conductive epoxy, 8329TCM, MG Chemicals) into the corresponding cut-out in the sample holder. For the closed-loop temperature control a thermistor (NTC thermistor, 10k, GA10K3MCD1), positioned within the sample holder as close as possible to the microscope dish, and a temperature controller (TC200, Thorlabs) was used. A spring-loaded design for the connection between the sample holder and the temperature controller (Supplementary Figure 6A) simplifies the assembly and handling of the sample chamber and improves the durability. By minimizing the contact surface area between the sample holder and the sample holder frame by a three-point contact system, we effectively reduce heat dissipation to the sample holder frame, which may act as a heat sink, ensuring temperature stability. For the fluid exchange microfluidic tubes (Elastosil R plus, 0.76 x 1.65 x 0.45 mm) are used. The waste media which is possibly contaminated is directed into a closed vessel containing disinfectant by creating negative pressure in the waste container via a syringe and a 0.2 µm filter in between.

### Custom AFM setup

AFM images were recorded using a custom AFM setup (Supplementary Figure 7). The AFM scan head (Dimension 3100, Digital Instruments) was mounted above an inverted optical microscope (IX81, Olympus). Illumination was provided by a mercury lamp (U-HGLGPS, Olympus) and fluorescence images were acquired with a sCMOS camera (Zyla 4.2 Plus, Andor). An open-source FPGA-based SPM controller (National Instruments USB-7856) and a corresponding LabVIEW-based software were used to control the AFM. A custom high-voltage amplifier was used to drive the piezoelectric actuators of the AFM scanner. The stepper motors for x-y-z translation of the AFM scan head and the sample chamber were controlled by micro-stepping analog drivers (M542, Leadshine). An additional interconnect module contains a custom printed circuit board (PCB), which interfaces the SPM controller and the AFM scan head and includes a display for facilitating laser alignment.

### Assessment of AFM performance

The noise level of the AFM setup was quantified under different sample chamber conditions. The complexity of the sample chamber assembly was incrementally increased to resemble the final assembly, and the root mean square (RMS) noise was measured throughout this process (Supplementary Figure 9A). Scans were performed in contact mode using a low gain setting with a scan rate of 1 Hz, 256 pixels per scan line (sampling rate: 1 Hz x 256 pixels/line = 512 pixel/s = 512 Hz (forward, backward), effective measurement bandwidth: 512 Hz / 2 = 256 Hz (Nyquist sampling theorem), 256 scan lines per frame, and a scan size of 1 nm x 1 nm. The data was acquired using ScanAsyst Fluid cantilevers (Bruker) with a nominal spring constant of 0.7 N m^-1^. The RMS noise over the measurement bandwidth was determined using the software Gwyddion and averaged over multiple measurements. The noise level in nanometers was calculated by multiplying the RMS noise by the deflection sensitivity, which was recorded using force-distance curve measurements.

A custom module of the LabView software was used to measure the frequency response of the piezoelectric z-actuator within the main structure. The module uses a pseudo-random binary sequence (PRBS) as the input signal for the z-piezo actuation and measures the deflection. The Fourier transform of the input-output data gives the frequency response corresponding to the system under study. The following parameters were used for the system identification: PRBS order 12, PRBS length 4095, sub-nanometer amplitude excitation. The frequency response was measured for different conditions in liquid (Supplementary Figure 9B). To evaluate the effect of the aperture diameter, the microscope dish was attached to a metal plate with circular cutouts of varying diameters (no cutout, cutout with ∅10 mm, cutout with ∅30 mm). For each condition, the microscope dish was positioned centric to the corresponding cutout. To evaluate the effect of the liquid volume, the microscope dish was placed in the sample holder (aperture ∅10 mm) and various liquid volumes (0.25 ml, 2 ml, 4 ml) were pipetted into the center of the microscope dish. The measurements were recorded in contact mode, at a sampling frequency of 125 kHz and averaged with 100 repetitions per condition. The Bode plots were plotted with MATLAB.

The simulation for the Eigenfrequency analysis (Supplementary Figure 9C) was performed with the finite element simulation software COMSOL Multiphysics® version 6.1. The CAD model for the sample holder assembly with microscope dish, was created using SOLIDWORKS. The materials used in the simulation were Aluminium 6063-T83, Schott N-BK7, Acrylic plastic.

### Microscope dish functionalization

*M. abscessus* (smooth and rough variant) and *M. marinum* bacteria were immobilized using Polydimethylsiloxane (PDMS). The microscope dishes were PDMS coated according to a modified protocol^10^. In short, the microscope dishes were rinsed with ethanol, dried with nitrogen and plasma-activated. PDMS was mixed in a ratio of 1:10, air-bubbles were removed by degassing under vacuum, a drop of PDMS was spin-coated at 8000 rpm for 45 s and the PDMS-coated microscope dish was baked at 60 °C for 4 h. The temperature during the baking step should not be set higher than 60°C to avoid deformations of the plastic part of the microscope dish, possibly leading to instabilities when mounting the microscope dish into the sample holder.

### Bacterial culture conditions

*M. abscessus* ATCC 19977 (smooth and rough variant, wild-type), *M. marinum M* (wild-type) and derivative strains were cultured in Middlebrook 7H9 liquid medium (Difco) supplemented with 0.5% albumin, 0.2% glucose, 0.085% NaCl, 0.5% glycerol, and 0.05% Tween-80. *M. abscessus* cultures were grown at 37 °C, while *M. marinum* cultures were grown at 30 °C, until they reached an optical density at 600 nm (OD 600 nm) of approximately 0.5, indicative of the mid-exponential phase. The *M. marinum* GFP reporter strain was grown in growth medium supplemented with kanamycin (50 μg ml^-1^). Aliquots were stored in 15% glycerol at −80 °C and thawed at room temperature before use. Each aliquot was used once and then discarded.

### *M. marinum* fluorescent reporter strain

A *M. marinum* reporter strain constitutively expressing GFP was constructed by transformation with the msp12::GFP plasmid. The plasmid expressing GFP has been described previously^38^. msp12::GFP was electroporated into *M. marinum* and resulting transformants were selected by plating on solid 7H9 medium containing 50 μg ml^-1^ kanamycin.

### Bacterial immobilization

Aliquots from exponentially phase cultures of *M. abscessus* (smooth and rough variant) and *M. marinum* were pipetted onto PDMS-coated coverslips, covered with a glass coverslip and incubated at 37 °C for 1 h to allow the cells to adhere by passive sedimentation. Unattached bacteria were removed by washing the microscope dish with 7H9^10^. *M. abscessus* rough variant and *M. marinum* were filtered before immobilization as they are more prone to form cell clumps. Cell culture aliquots were centrifuged (5000 rpm, 5 min), the supernatant was removed and the cell pellet was resuspended in 7H9. Cells were filtered through a 5 µm pore size PVDF filter (Millipore).

### Fluorescent staining

M. abscessus (smooth variant) was stained with BODIPY™ FL Vancomycin (life technologies, V34850) with a protocol adapted from Thanky et al.^39^. Bacteria were grown to mid-exponential phase, collected by centrifugation (5000 rpm, 5 min) and incubated with 1 mg/ml BODIPY™ FL Vancomycin for 5h in a shaking incubator at 37 °C. Cells were centrifuged to remove excess stain and resuspended in 7H9 liquid medium. After the washing step was repeated twice, the cells were immobilized on PDMS as described above.

### Image acquisition

AFM images were acquired using ScanAsyst Fluid cantilevers (Bruker) with a nominal spring constant of 0.7 N m^-1^ in off-resonance tapping (ORT) mode^40^ at an oscillation rate of 500 - 1 kHz, force setpoints <3 nN at line rates of 0.2-0.5 Hz. Fluorescence images (pixel format 1024 × 1024) were recorded with a 100x objective (1.3 NA, oil immersion). The AFM laser was turned off during fluorescence image acquisition. The samples were maintained at 37 °C (*M. abscessus*) or 30°C (*M. marinum*) in 7H9 medium without Tween-80. The values of the stiffness channel are based on the slope of the approach curve. To calculate the approach slope, the deflection and the cantilever motion were integrated between the manually selected minimum deflection point and maximum deflection point.

### Image processing

AFM images were processed with the scanning probe visualization and processing software Gwyddion (Gwyddion). The color range was adjusted for all image channels. For the topography image, rows were aligned, scars were corrected for and the image was leveled. The line profiles of the stiffness channel were measured with Gwyddion and visualized with MATLAB. Outliers in the stiffness channel were removed using a custom Python script. A threshold was manually defined based on the histogram, ensuring it was set within the 0.95 to 0.99 quantile range. This threshold was then applied for data visualization to clip any values exceeding it. ImageJ was used to process the fluorescence images. The fluorescence images were cropped to the corresponding AFM scan size and a background subtraction was performed. The fluorescence image of the *M. marinum* reporter strain was filtered with a Gaussian filter (σ = 2). If an AFM scan angle ≠ 0 ° was used, the fluorescent image was rotated accordingly by bilinear interpolation.

## Data availability

Data for this article, including Supplementary Videos are available at a Zenodo repository at https://zenodo.org/records/13593196. Additional data supporting this article have been included as part of the Supplementary Information.

## Acknowledgments

This research was funded by the EPFL Open Science Fund (Open hardware workshops, 7607; G.E.F), the European Research Council (ERC-2017-CoG, InCell, 773091; G.E.F), Innosuisse (Eurostars-Eureka-AAL-ECSEL, E!665, 9953; NAFTAQ: Novel AFM Techniques for Autonomous Quality Control in Industrial Manufacturing, 102.271 IP-EE, 10632; G.E.F), the ETH Domain ORD Program (Open SPM, 10126; G.E.F), the SNSF (Real-Time Single-Cell Analysis of Bacterial Persistence, 310030_207717; J.D.M.). The authors want to acknowledge O. Rutschmann for constructing the GFP - *M. marinum* reporter strain.

## Author Contribution Statement

E.B., J.D.M. and G.E.F. conceptualized the idea. E.R. designed, developed, manufactured the sample chamber and performed the functional testing. E.R. constructed and assembled the custom combined optical and AFM setup. S.H.A, M.K., N.S.A. contributed to the development of the sub-systems of the custom AFM setup and the used custom control software, supported the assembly and testing of the sub-systems and provided advice for the project. E.B designed, conducted the experiments and collected, processed, analyzed the data, wrote and edited the manuscript. J. M. and G.E.F supervised and provided funding. All authors reviewed, edited and approved the manuscript

