## Supplementary Information for "A hermetically closed sample chamber enables time-lapse nano-characterization of pathogenic microorganisms *in-vitro*"

[*] Corresponding author

#### Supplementary Information:

**Assessment of AFM functionality and imaging performance**

To evaluate the AFM imaging performance, we first assessed the noise level under different conditions. In order to do so, we measured the noise level while gradually increasing the complexity of the sample chamber, indicating the resemblance with the final condition. We subsequently added the individual components of the sample chamber in a systematic order. Thereby, we attempted to detangle the possible effect of the different subcomponents of the sample chamber to evaluate their effect on the mechanical noise level. As a baseline condition for comparison, we used a commercial cantilever holder (Bruker, Dimension cantilever holder fluid) for AFM imaging in liquid and measured directly on a fixed metal plate without employing the sample holder. All measurements, including the baseline condition, were acquired with a custom AFM setup. We show that the complete sample chamber assembly does not lead to higher mechanical noise levels up to a measurement bandwidth of 256 Hz relative to the described baseline condition, both measured with the same custom microscope setup. The customized cantilever holder does not have a higher noise level compared to the commercial cantilever (quantitative values in Supplementary Figure 8A). In addition to employing the inner clamp, rubber O-ring, and vacuum grease for sealing, these components also contribute to mechanically stabilizing the microscope dish, thereby improving the noise level. By optimizing the size of the outer diameter of the rubber O-ring, we were able to further reduce the noise level. The O-ring is positioned within a circular cutout at the base of the sample holder, and its outer diameter determines how much of it protrudes. Our objective was to achieve an optimal configuration where, upon fastening the microscope dish, contact between the microscope dish and both the sample holder and the O-ring is ensured. A large contact area between the microscope dish and sample holder enhances the rigidity and the contact with the O-ring can support dampening, thereby enhancing the overall mechanical stability. The use of the membrane along with its support structure also helps reduce the noise level, likely by providing additional stabilization and dampening, as the flexible membrane material can absorb and dissipate vibrations. However, we also identified sources of mechanical noise. The noise level increases with a larger mechanically unsupported area of the microscope dish. An increased unsupported area reduces the contact interface between the sample holder and the microscope dish, which compromises structural stability and creates a more compliant mechanical system. As a result, external mechanical vibrations are more easily transmitted, increasing the system's susceptibility to disturbances and potentially elevating the noise level. Furthermore, larger liquid volumes within the microscope dish impact the noise level slightly detrimental due to the increased overall mass and higher hydrodynamic coupling between the z-scanner motion and the microscope dish.

We also characterized the effect of the microscope dish on the frequency bandwidth of the system (i.e. z-actuator and microscope dish coupled via liquid) by measuring the frequency response of the system under different conditions (Supplementary Figure 8B). In our study, we observed that a mechanically unsupported area (defined by the aperture size) of the sample holder and the use of liquid (e.g. growth media) within the sample holder generate dynamics that interfere with the feedback loop by reducing the phase and gain margins, therefore affecting the system’s bandwidth (i.e. usable frequency range) and limiting the scan rate. The larger the aperture diameter and the larger the media volume, the lower the frequency of these dynamics, resulting in a smaller usable frequency range and a lower possible scan rate. We found a trade-off between the aperture size, which affects the potential addressable region of interest for optical imaging, with the need for a large bandwidth to achieve effective correlated imaging.

We also performed an Eigenfrequency analysis of the sample holder with microscope dish and inner clamp and observed that the mechanically unsupported area is more prone to vibrate and that the eigenmode deformation is most pronounced in the center of the unsupported area (Supplementary Figure 8C).

#### Supplementary Figures:

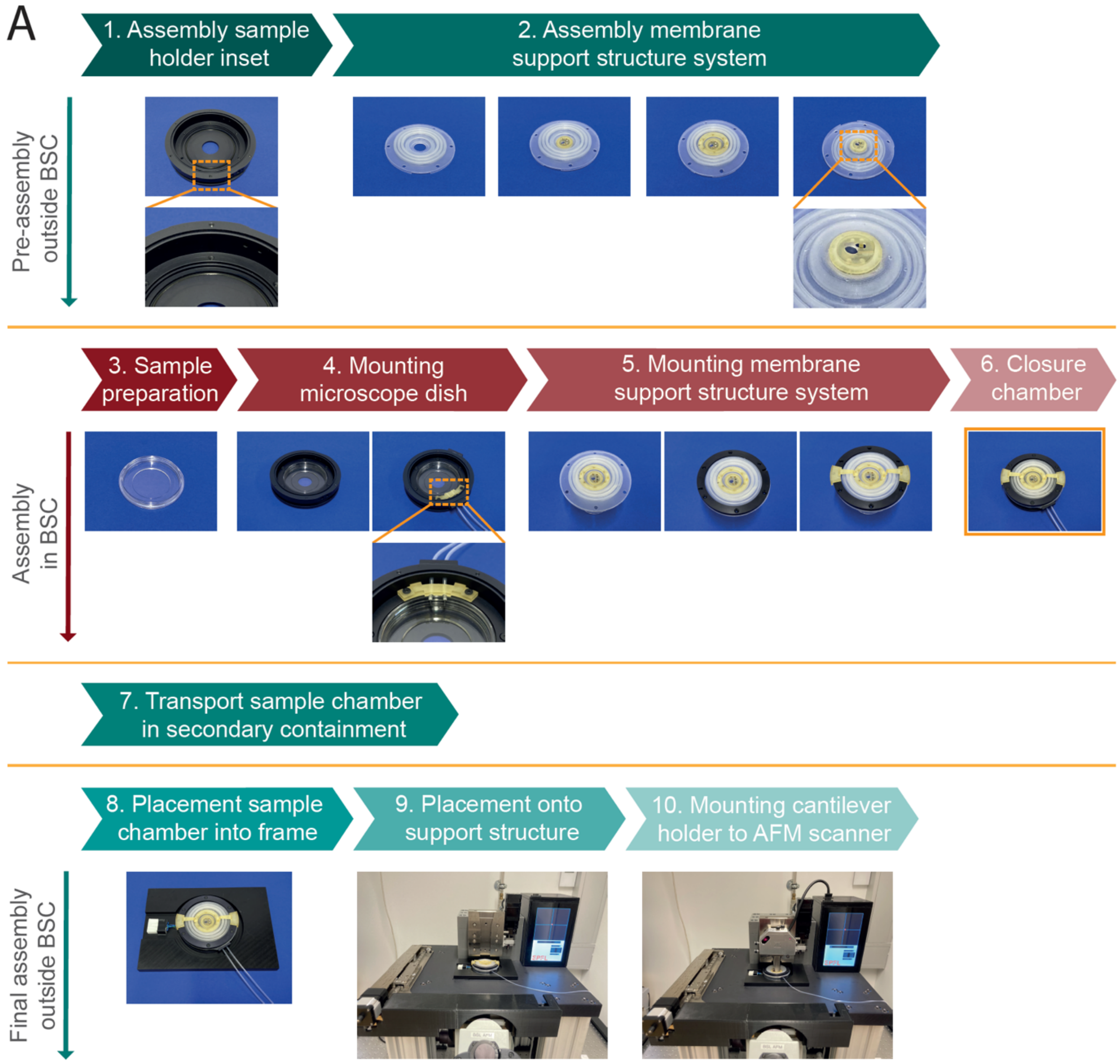

**Supplementary Figure 1: Workflow of the sample chamber assembly procedure**

Pre-assembly outside of a biosafety cabinet (BSC): (1) The rubber O-ring is placed into the corresponding cut-out at the bottom of the sample holder. (2) The cantilever holder and the membrane support structure are mounted to the membrane. Afterwards, the cantilever is mounted to the cantilever holder. Assembly in a BSC: (3) Sample immobilization and corresponding washing steps are performed in the microscope dish. (4) The microscope dish is placed into the sample holder and fastened with the inner clamps. The tubes are inserted into the fluid exchange ports and fixed with the tube fixation. The fluid exchange ports are sealed with O-rings and the outer clamps. (5) The outer clamp is placed onto the membrane and the assembly support structure is slit into the outer clamp to fixate the cantilever holder position. (6) The membrane with membrane support structure system is placed on top of the sample holder and fastened. (7) The closed sample chamber is transported to the

AFM setup in a secondary containment. Final assembly outside BSC: (8) The sample chamber is placed into the frame. (9) The frame with the sample chamber is placed onto the AFM support structure above the inverted optical microscope. (10) The AFM scanner is placed above the sample chamber and the AFM scanner is attached to the cantilever holder.

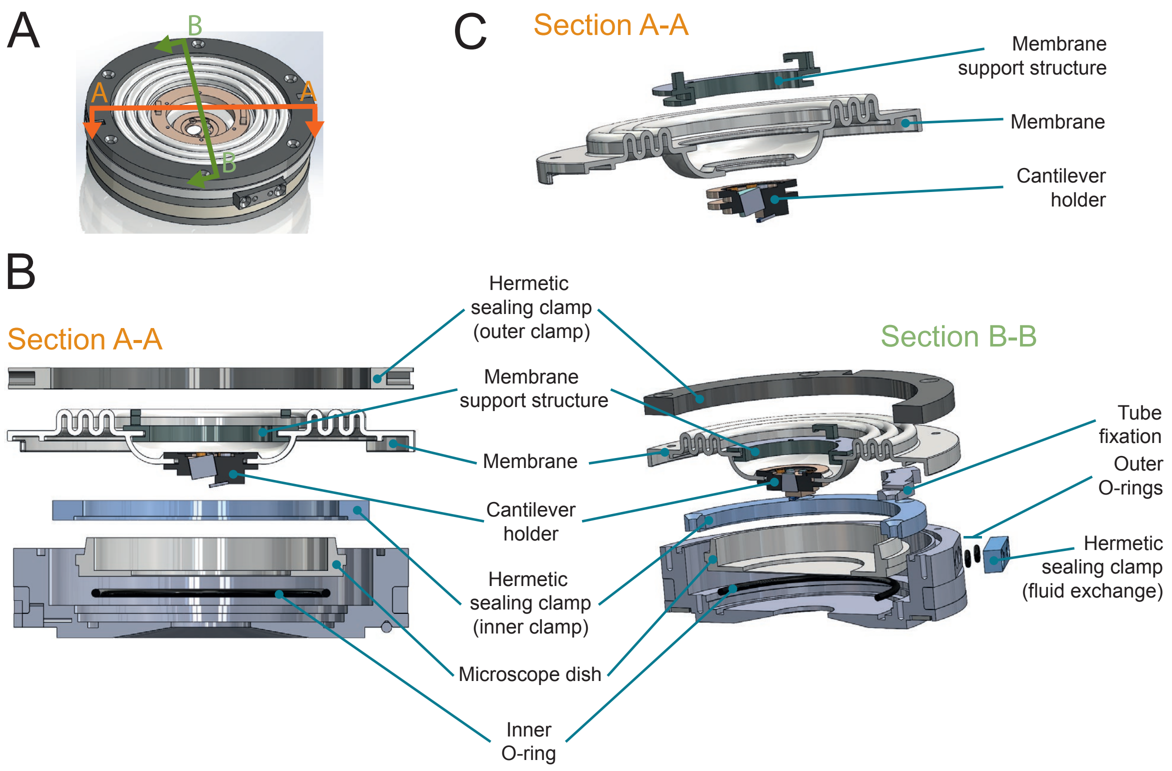

**Supplementary Figure 2: Explosion views of the hermetically sealed AFM sample chamber with membrane support structure**

**A** 3D rendering of the sample chamber with lines indicating the cross-sections and direction of view used for the explosion views in the sub-panels. **B** Explosion views of cross-sections A-A and B-B with corresponding sample chamber components. **C** Explosion view of cross-section A-A depicting the components mounted to the membrane via an interference fit.

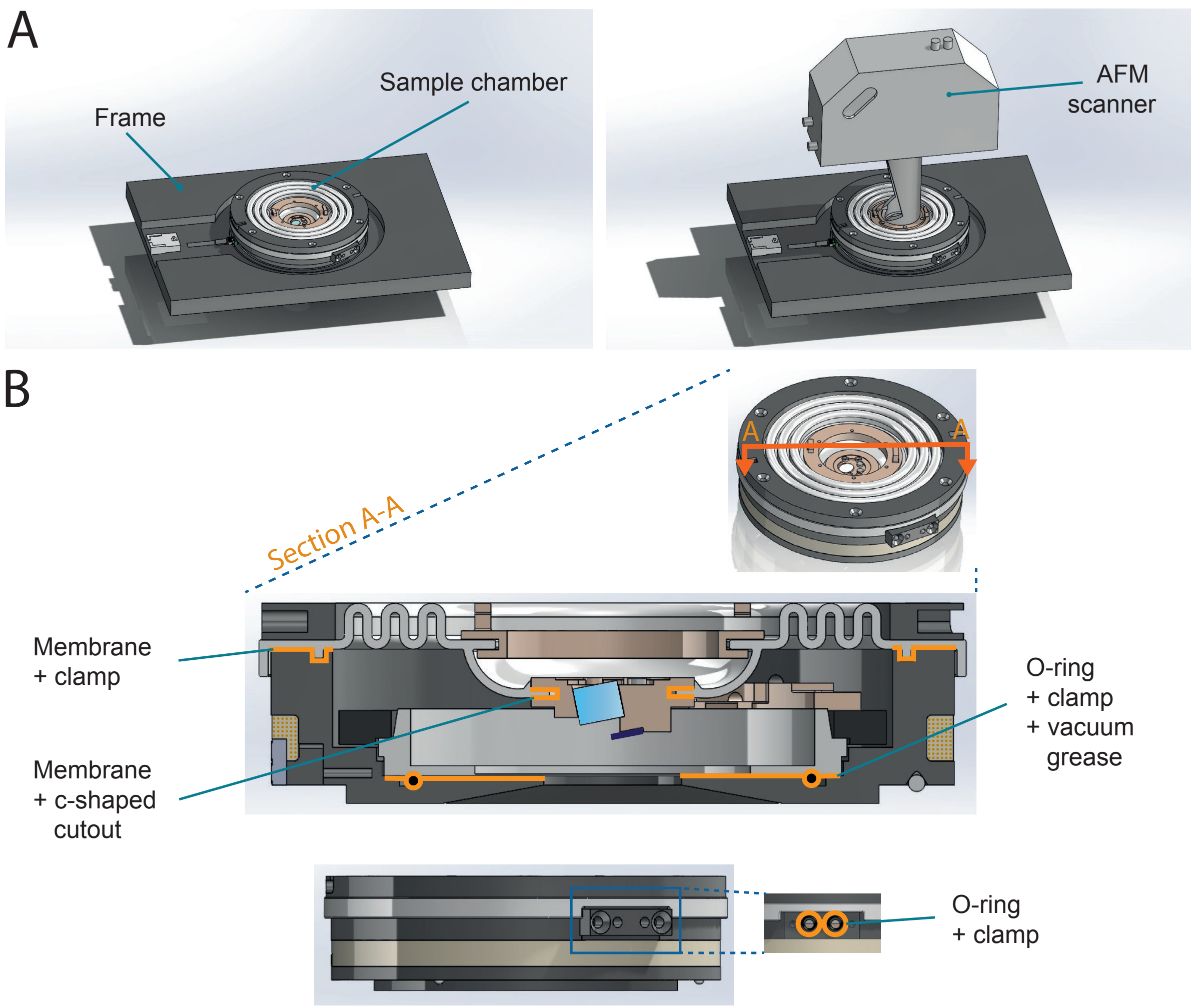

**Supplementary Figure 3: Sample chamber and corresponding sealing points**

**A** 3D rendering of the complete sample chamber before (left) and after (left) mounting it to the AFM scanner. To improve the mechanical stability and handling during imaging, the sample chamber is placed into a frame via a magnetic kinematic mount. **B** 3D rendering of sample chamber and cross-section. The multiple sealing points ensuring hermetic sealing are highlighted in orange. The chamber is closed from the top by a specially designed silicone membrane, which is secured at the outer edge to the sample holder by a clamp and at the inner edge to the cantilever holder via an interference fit. The interface at the bottom of the sample holder is sealed by a combination of an O-ring and vacuum grease placed underneath the microscope dish and a clamp. The fluid exchange ports are sealed with O-rings and clamps.

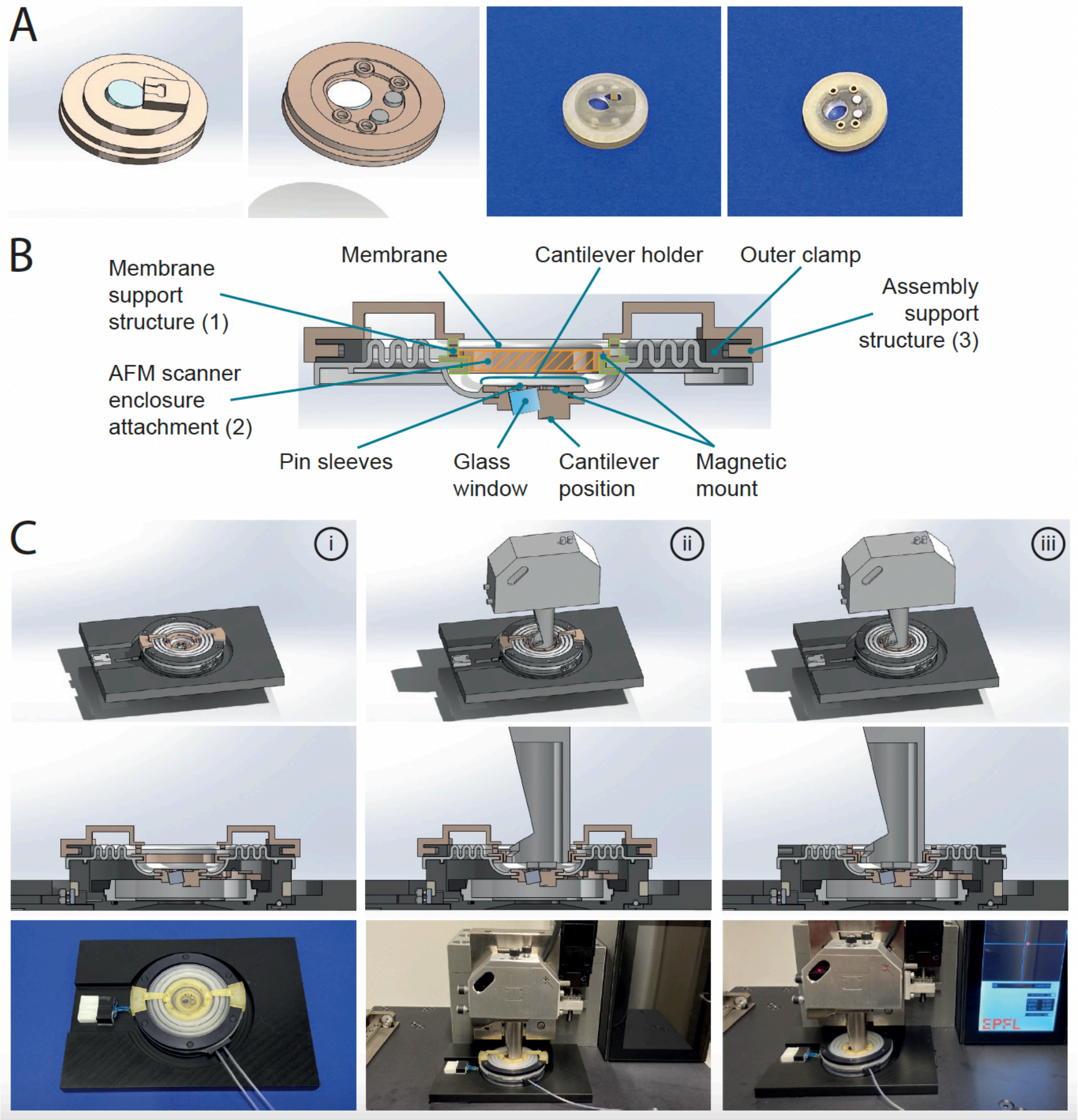

**Supplementary Figure 4: Cantilever holder and membrane support structure system**

**A** 3D rendering (left) and images (right) of the cantilever holder. To enable mounting the membrane to the cantilever holder by an interference fit, the cantilever holder features a specifically designed c-shaped cutout. **B** Cross-section of 3D rendering of the membrane support structure system with membrane and cantilever holder. This system consists of three sub-components (1-3). A circular support structure (1) [membrane support structure] is attached to the membrane by an interference fit. The membrane support structure is also attached to an anchor point permanently fixed to the AFM scanner enclosure (2) [AFM scanner enclosure attachment] via a magnetic mount. In this way, the membrane support structure holds a significant portion of the membrane's weight via this additional anchor point. Additionally, support arms (3) [assembly support structure] can be temporarily attached to the membrane support structure and outer clamp via a slot-and-tab mount, facilitating the transport of the sample chamber while preventing the cantilever holder from descending, thus protecting the fragile cantilever from potential damage. The membrane support structure (green) and the AFM scanner enclosure attachment (orange) are outlined for better differentiation. The cross-hatched area indicates the surface of the AFM scanner enclosure attachment, which is permanently fixed to the scanner enclosure. **C** 3D rendering of sample chamber with cross-section (top rows) and images (bottom row) of sample chamber and AFM scanner assembly. (i) During sample chamber assembly and transport to the AFM setup, the assembly support structure is mounted to the sample chamber. (ii) The AFM scanner is attached to the cantilever holder via a magnetic mount. (iii) After the cantilever holder is attached to the AFM scanner, the assembly support structure is removed by sliding it out of the slot-and-tab mount.

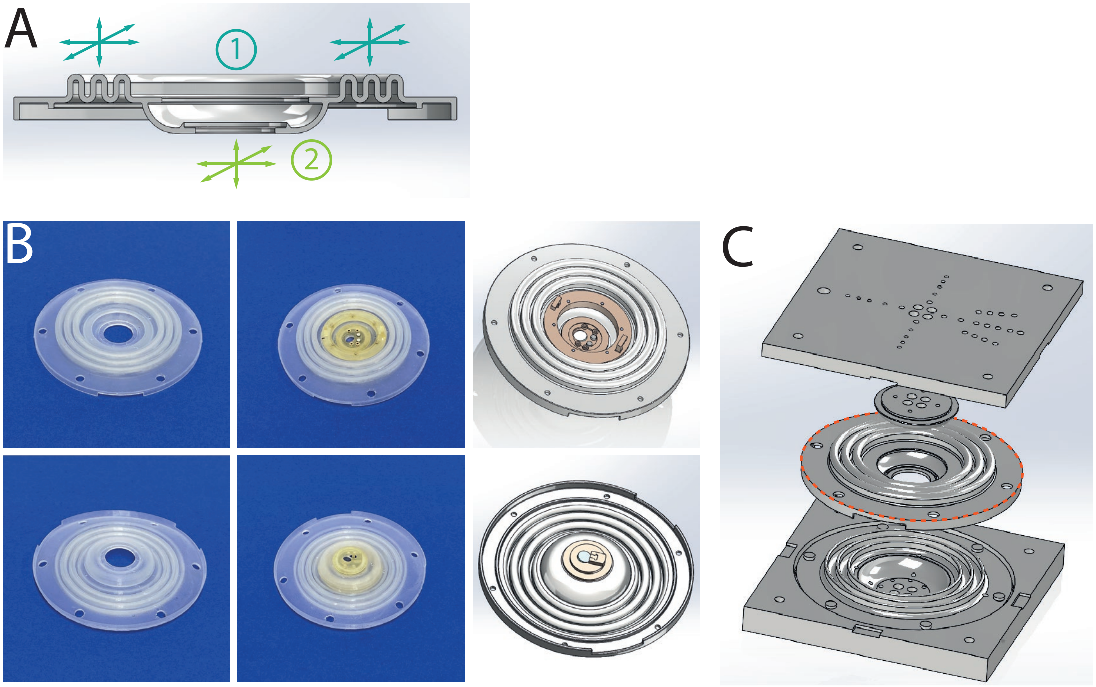

**Supplementary Figure 5: Membrane design and membrane mold**

**A** Cross-section of 3D rendering of the membrane with indicated direction of flexibility. The membrane is designed for flexibility on two levels. (1) The outer part permits x-y-z movement of the sample chamber relative to the AFM scanner housing. (2) The inner part allows x-y-z clearance for the AFM scanner movement relative to its housing. **B** Images of the silicone membrane (left) and membrane with attached cantilever holder and membrane support structure (middle). 3D rendering of membrane with cantilever holder and membrane support structure (right). **C** 3D rendering of the membrane mold for silicone injection molding and the resulting membrane (orange dotted line).

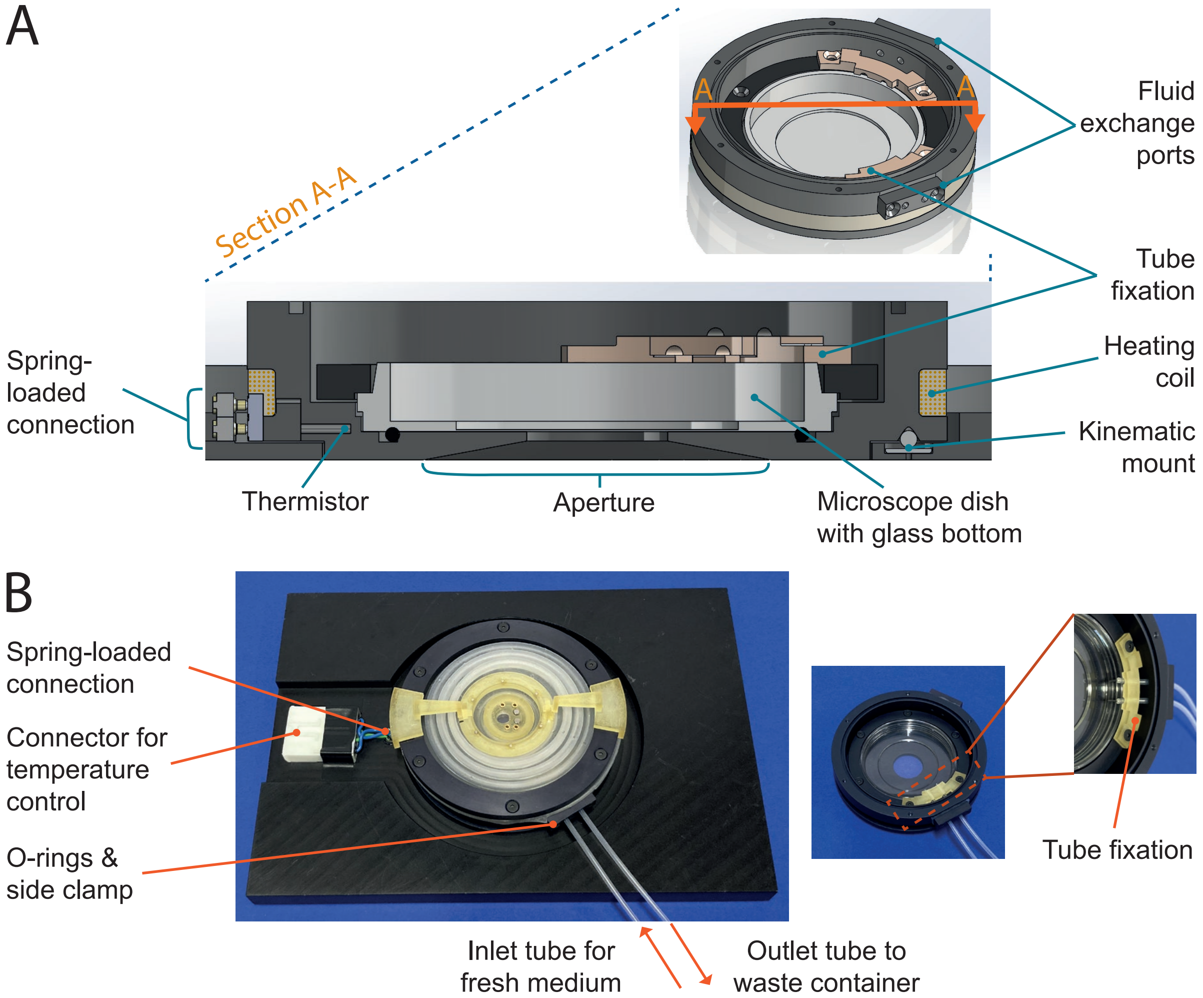

**Supplementary Figure 6: Additional sample chamber features enabling time-lapse correlated optical and AFM imaging**

**A** Cross-section (A-A) of the 3D rendered sample chamber. Temperature control is implemented by using a heating coil as the heating element and a thermistor as the temperature sensor. A spring-loaded connector facilitates the connection between the heating element and the temperature controller. A kinematic mount serves as the interface between the sample holder and the frame. The aperture at the bottom of the sample holder allows objective access to the microscope dish. Fluid exchange is enabled through tubes inserted into the fluid exchange ports, which are secured using the tube fixation clamp. **B** Image of sample chamber showing the components for temperature and fluid control.

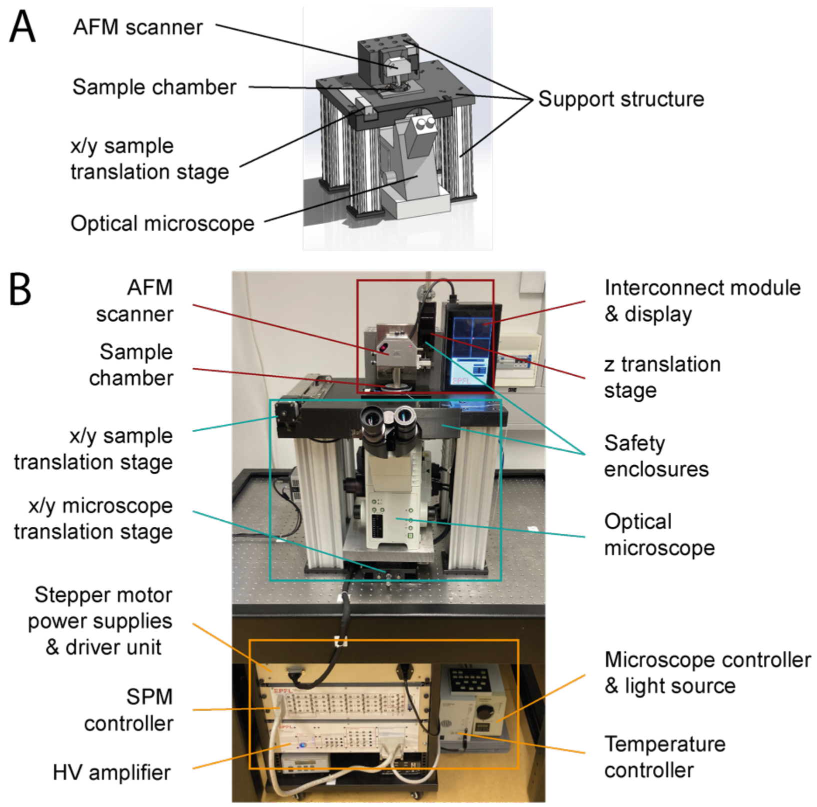

**Supplementary Figure 7: Complete combined optical and AFM setup**

**A** 3D rendering of the combined optical and AFM microscope setup showing the AFM scanner, sample chamber, inverted optical microscope and x/y sample translation stage. The support structure facilitates the positioning of the sample chamber above the inverted microscope and the AFM scanner above the sample chamber while ensuring mechanical stability. The complete microscope is placed on an anti-vibration table. **B** Image depicting the different sub-systems of the setup with the individual components.

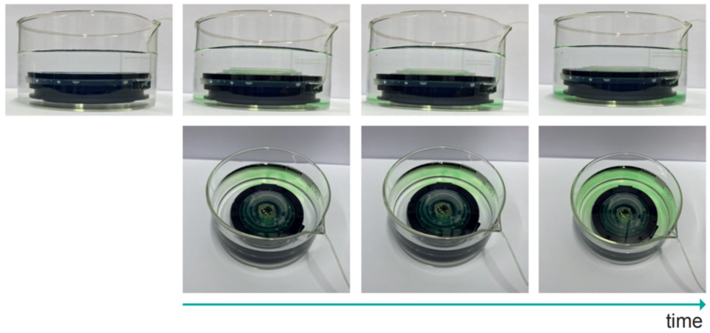

**Supplementary Figure 8: Control leakproofness test with pinched membrane**

Timelapse images of leakproofness test with pinched membrane. The sample chamber with pinched membrane was submerged in transparent liquid and then pressurized with dyed liquid. Over time, a discoloration of the surrounding media due to the leakage of dyed liquid was observed.

**A**

| Conditions | mean RMS noise (nm) |
|---|---|
| commercial cantilever holder | 0.18 |
| without membrane & membrane support structure | 0.16 |
| with membrane & membrane support structure | 0.20 |
| o-ring & clamp - without membrane & membrane support structure | 0.18 |
| o-ring & clamp - with membrane & membrane support structure | 0.12 |
| without o-ring & clamp - with membrane & membrane support structure | 0.25 |
| without o-ring - with clamp - with membrane & membrane support structure | 0.25 |
| with o-ring & clamp - with membrane & membrane support structure | 0.08 |
| 2 ml - with o-ring & clamp - with membrane & membrane support structure | 0.11 |
| 4 ml - with o-ring & clamp - with membrane & membrane support structure | 0.18 |

metal plate

BSL sample holder

BSL cantilever holder

0.25 ml liquid

Complexity of assembly - Resemblance final assembly

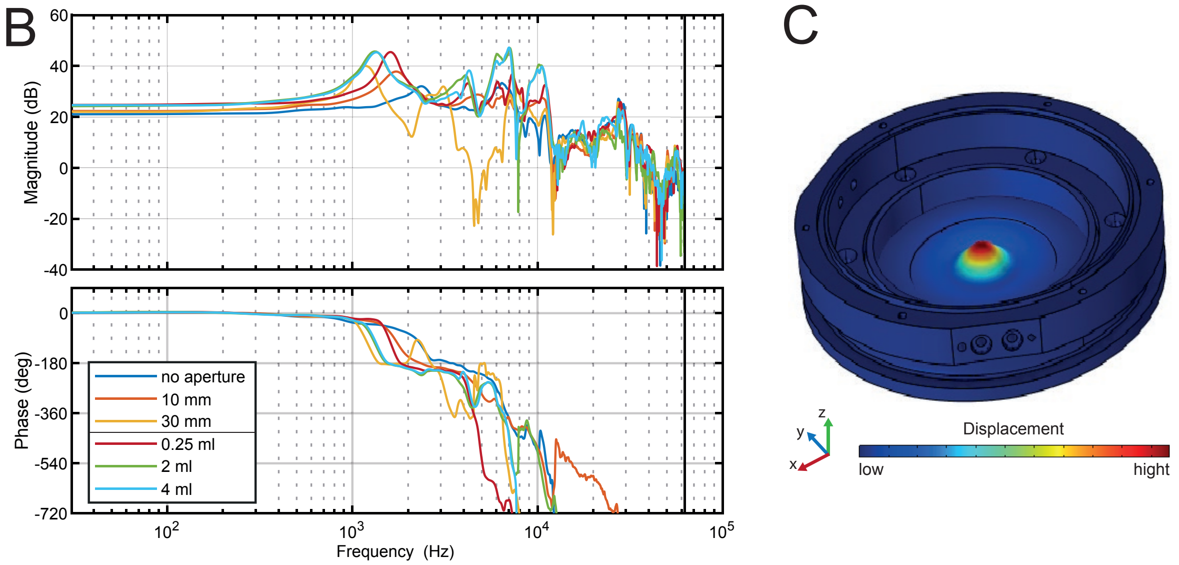

**Supplementary Figure 9: Evaluation of AFM functionality and imaging performance**

**A** Table with noise levels under different conditions. The complexity of the sample chamber, indicating the resemblance with the final assembly, was increased while measuring the mean RMS noise at 256 Hz measurement bandwidth. As a baseline condition for comparison, we used a commercial cantilever holder and measured directly on a fixed metal plate without employing the sample holder. All measurements, including the baseline condition, were acquired with a custom AFM setup. **B** Bode plot of the frequency response (i.e. cantilever deflection as a response to z-actuator excitation) of the system (i.e. z-actuator and microscope dish coupled via liquid) under different conditions. Larger aperture diameters of the sample holder and larger liquid volumes decrease the frequency of the generated dynamics limiting the system's frequency bandwidth. **C** Mode-shape representation of the first eigenmode of the sample holder with microscope dish and inner clamp. The most pronounced displacement is in the center of the microscope dish within the area of the sample holder aperture.

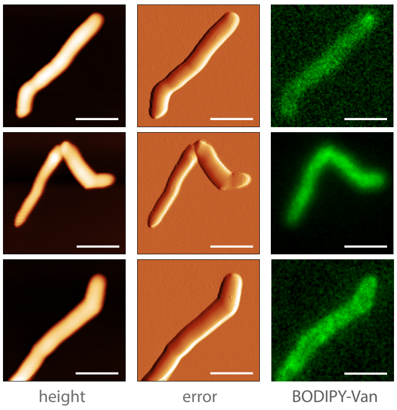

**Supplementary Figure 10: Combined optical and AFM imaging**

Correlated optical and AFM image of *M. abscessus* smooth variant stained with BODIPY™ FL Vancomycin (scale bar 2µm).

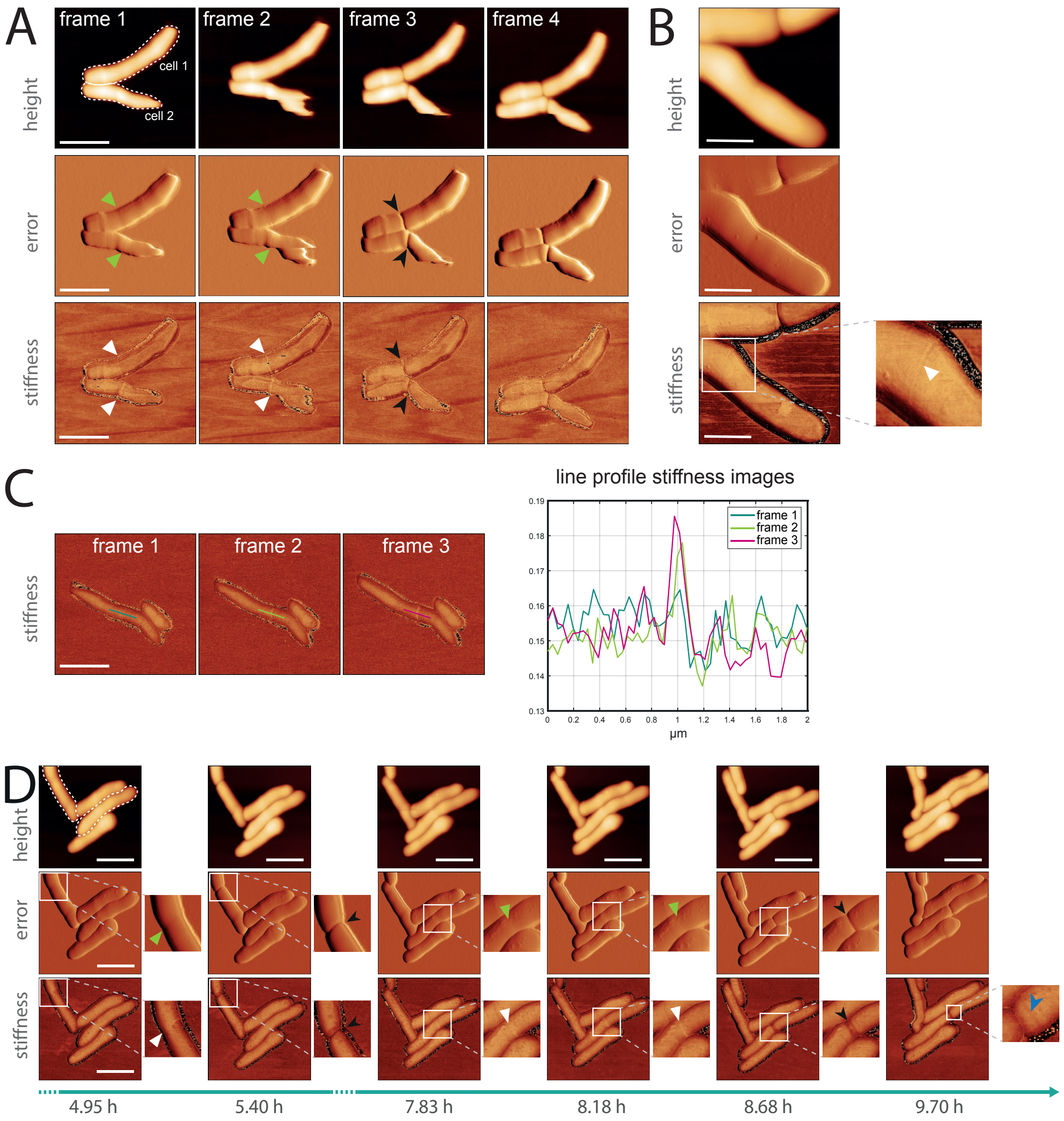

**Supplementary Figure 11: Characterization of cell cycle events of *M. abscessus* smooth variant (BSL 2)**

**A** Timelapse AFM images of *M. abscessus* cells (smooth variant) before and after cell cleavage. The images show the formation of the pre-cleavage furrow (green arrowhead) and an elevated level of stiffness in the area surrounding the pre-cleavage furrow (white arrowhead), which increases over time (frames 1, 2). Cell cleavage occurs at the location of the pre-cleavage furrow (black arrowhead, frame 3). The tip of cell two on the right was not fully immobilized on the surface, resulting in slight movement during AFM imaging. A v-snapping motion, which is a characteristic phenotype of the mycobacterial cell cleavage, may have amplified this movement. (scale bar 2µm). **B** AFM image of an *M. abscessus* cell (smooth variant) with a detailed view (digital zoom-in) showing an emerging pre-cleavage furrow and the associated increased stiffness (white arrowhead) (scale bar 2 µm). **C** Timelapse AFM images of an *M. abscessus* cell (smooth variant) before cell cleavage, showing the formation of the pre-cleavage furrow with elevated stiffness (left) and plotted line profiles in the stiffness channel drawn over the pre-cleavage furrow (right). The stiffness at the PCF increases over time (frames 1-3) (scale bar 1 µm). **D** Timelapse AFM images of *M. abscessus* cells (smooth variant) with detailed views (digital zoom-in). Two cells (dotted outline) display the formation of the pre-cleavage furrow (green arrowhead) along with an increase in stiffness (white arrowhead). Cell cleavage occurs at the site of the pre-cleavage furrow (black arrowhead). Following cell cleavage, a division scar becomes visible (blue arrowhead) (scale bar 2 µm).

### Supplementary Videos:

Supplementary videos are uploaded to a Zenodo repository

- **Video 1: Airtightness control** Control of airtightness with pinched membrane. The sample chamber was assembled with a pinched membrane, inflated with air and monitored for deflation.
- **Video 2: Airtightness control submerged in water** Control of airtightness with pinched membrane. The sample chamber was assembled with a pinched membrane, submerged in water, inflated with air and monitored for appearance of air-bubbles.
- **Video 3: Airtightness test** The sample chamber was assembled, inflated with air and monitored for deflation over a 24 h period.
- **Video 4: Airtightness test submerged in water** The sample chamber was assembled, submerged in water, inflated with air and monitored for appearance of air-bubbles over a 1 h period.
- **Video 5: Leakproofness test** The sample chamber was assembled, pressurized with a dyed liquid, submerged in a transparent liquid and monitored for maintenance of transparency over a 24 h period.

### Supplementary Tables:

#### Supplementary Table 1: List of used materials and decontamination options

| Material | Component | Decontamination by disinfectant* | Decontamination by autoclaving | Temperature range |
|---|---|---|---|---|
| 7075 aluminium alloy, anodized | Sample holder | chemical resistance | ✓ | to 477 °C |
| 6082 aluminium alloy, anodized | Plate of AFM support structure | chemical resistance | ✓ | to 555 °C |
| SF13 RTV-2 silicone | Membrane | chemical resistance | ✓ | - 40 - 200 °C |
| FPM | O-ring | chemical resistance | ✓ | - 20 - 200 °C |
| PC-like translucent (Accura 5530) | Membrane support structure, cantilever holder | not specified, stable based on experience | ✓ | to 170 °C - 250 °C |
| 8329TCS epoxy | Heat conductive glue | not specified, stable based on experience | ✓ | - 65 - 150 °C |
| 9200 epoxy | Structural epoxy | chemical resistance | ✓ | - 40 - 150°C |
| N-BK7 (coated) | Window cantilever holder | N-BK7 chemical resistance | ✓ | N-BK7 to 160°C |

* Depending on the pathogen (ethanol, Incidin, hydrogen peroxide)

#### Supplementary Table 2: Protocol to test general leakproofness and airtightness of sample chamber and while simulating multiple imaging sessions and membrane aging

| | |
|---|---|
| **1. Leakproofness assessment (general)** | |
| 1.1 | Closing sealed sample chamber at atmospheric pressure |
| 1.2 | Pressurization sample chamber with a dyed liquid (~20 ml) |
| 1.3 | Turning and shaking closed sample chamber and testing sealing points for leakage of fluid |
| 1.4 | Submerging sample chamber in container with transparent liquid |
| 1.5 | Monitoring the transparency over a period of 24 h |
| **2. Airtightness assessment in air (general)** | |
| 2.1 | Closing sealed sample chamber at atmospheric pressure |
| 2.2 | Pressurization sample chamber with air (~500 mBar) leading to a bulging membrane |
| 2.3 | Monitoring for deflation of the bulging membrane of over a period of 24 h |
| **3. Airtightness assessment in liquid (general)** | |
| 3.1 | Closing sealed sample chamber at atmospheric pressure |
| 3.2 | Pressurization sample chamber with air (~500 mBar) leading to a bulging membrane |
| 3.3 | Monitoring for appearance of air-bubbles of over a period of 24 h |
| **4. Airtightness assessment in air (simulating multiple imaging sessions)** | |
| 4.1 | Closing sealed sample chamber at atmospheric pressure |
| 4.2 | Pressurization sample chamber with air (~500 mBar) leading to a bulging membrane |
| 4.3 | Depressurising to atmospheric pressure |
| 4.4 | Repeating step 4.2 and 4.3 over 100 cycles |
| 4.5 | Pressurization sample chamber with air (~500 mBar) leading to a bulging membrane |
| 4.6 | Monitoring for deflation of the bulging membrane of over a period of 24 h |
| **5. Airtightness assessment in air (simulating membrane aging)** | |
| 5.1 | Autoclaving membrane 20 times |
| 5.2 | Subsequently performing airtightness assessment in air (2.) |
